# Odd-chain dicarboxylic acid feeding recapitulates the biochemical phenotype of glutaric aciduria type 1 in mice

**DOI:** 10.1101/2025.02.13.637994

**Authors:** Adam C. Richert, Yuxun Zhang, Sivakama S. Bharathi, Abigail Hernandez, Tetyana Dodatko, Joanna Bons, Brandon Stauffer, Chunli Yu, Birgit Schilling, Sander M. Houten, Eric S. Goetzman

## Abstract

Glutaric aciduria type-1 (GA1) is an inherited mitochondrial neurometabolic disorder with a poorly understood pathogenesis and unmet medical needs. GA1 can be diagnosed via its hallmark biochemical signature consisting of glutaric aciduria, 3-hydroxyglutaric aciduria, and increased plasma glutarylcarnitine. These glutaryl-CoA-derived metabolites are thought to originate solely in the mitochondria. Here, we demonstrate that wild-type mice fed an 11-carbon odd-chain dicarboxylic acid (undecanedioic acid, DC_11_) recreates the biochemical phenotype of GA1. Odd-chain dicarboxylic acids like DC_11_ are not present in food but can arise from several endogenous processes, such as lipid peroxidation and fatty acid ω-oxidation. DC_11_ is chain-shortened in peroxisomes to glutaryl (DC_5_)-CoA, which then gives rise to the GA1-like pattern of DC_5_ metabolites in urine, tissues, and blood. Glutaric acid released from peroxisomes during DC_11_ chain-shortening can enter mitochondria, be activated to CoA by the enzyme succinyl-CoA:glutarate-CoA transferase (SUGCT), and become substrate for glutaryl-CoA dehydrogenase (GCDH), the enzyme that is mutated in GA1. Our data provide proof-of-concept that the generation of dicarboxylic acids by ω-oxidation, which is stimulated during the same catabolic states known to trigger acute encephalopathy in GA1, may exacerbate disease by increasing the glutaryl-CoA substrate load in mitochondria.

## INTRODUCTION

Glutaric aciduria type 1 (GA1) is an autosomal recessive inborn error of lysine, hydroxylysine and tryptophan degradation first reported in 1975 (MIM 231670)[1]. GA1 is rare (∼1 in 100,000 births), occurring most frequently in the Amish, Ojibwe and Lumbee Indigenous peoples, Irish Traveller Community, and Xhosa people of South Africa [2–4]. If left untreated, patients typically suffer from acute encephalopathic crises, leading to striatal injury and a complex movement disorder. The encephalopathic crises are often triggered by a catabolic state such as those that occur during childhood illnesses [5, 6]. GA1 patients benefit from immediate intervention and the disorder is therefore included in newborn screening programs [7, 8]. Current treatment consists of strict limitation of dietary lysine, along with carnitine supplementation and emergency interventions [9]. Even when the treatment regimen is meticulously maintained, neurological disease still develops in ∼25% of patients [7]. As early detection improves lifespan, long-term complications such as cognitive impairment, extra striatal neurotoxicity, subdural hematomas, and kidney disease are becoming more prominent and may require life-long management [10–12]. These limitations demonstrate the unmet need for additional novel therapeutic approaches.

GA1 is caused by mutations in the mitochondrial enzyme glutaryl-CoA dehydrogenase (GCDH) [13]. GCDH is a member of the acyl-CoA dehydrogenase (ACAD) enzyme family. There are nine ACADs with known functions and at least two additional enzymes that remain largely uncharacterized [14]. All ACADs dehydrogenate acyl-CoA substrates, effectively inserting a double-bond between the second and third carbons. GCDH is unique among ACADs in that it is also a decarboxylase, converting its five-carbon substrate glutaryl-CoA into a four-carbon product, crotonyl-CoA, plus CO_2_ [15, 16]. When GCDH is deficient, glutaryl-CoA accumulates inside mitochondria. Glutaryl-CoA cannot cross the mitochondrial membrane. As it accumulates, it becomes subject to three fates that produce the hallmark biochemical phenotype of GA1. First, glutaryl-CoA can be cleaved to glutaric acid and free CoA. Glutaric acid can cross biological membranes and leave the cell. Second, carnitine acyltransferase enzymes can exchange the CoA for carnitine, yielding glutarylcarnitine, which can also cross membranes. Third, at high enough concentrations glutaryl-CoA becomes a substrate for another ACAD enzyme, medium-chain acyl-CoA dehydrogenase (MCAD) [17]. MCAD dehydrogenates glutaryl-CoA, but since it lacks the decarboxylase function, MCAD produces an unsaturated five carbon intermediate known as glutaconyl-CoA rather than crotonyl-CoA. Glutaconyl-CoA can be hydrated by 3-methylglutaconyl-CoA hydratase (3MGH) to yield 3-hydroxyglutaryl-CoA [18]. These CoA esters hydrolyze to glutaconic and 3-hydroglutaric acid, respectively. Diagnosis of GA1 can be made on the presence of 3-hydroxyglutaric and glutaric acid in urine, along with glutarylcarnitine in blood. Accumulating metabolites, in particular 3-hydroxyglutaric and glutaric acid, are thought to contribute to striatal damage in GA1 [19, 20].

While glutaric acid is formed during degradation of an amino acid, chemically it belongs to a class of fatty acids known as dicarboxylic acids (DCAs). Whereas the fatty acids in our diet and stored as triglycerides in our bodies are “monocarboxylic”, having a carboxyl group on the α-carbon, DCAs also have a carboxyl group on the opposite (ω) end. DCAs are rare in food. Rather, they are synthesized from monocarboxylic fatty acids in the liver and kidney through a minor pathway known as ω-oxidation. Conversion of monocarboxylic acids into DCAs begins with a hydroxylation step catalyzed by cytochrome P450 enzymes in the endoplasmic reticulum followed by two oxidation steps in the cytoplasm [21]. It is thought that the resulting DCAs are preferentially β-oxidized by the peroxisomal fatty acid oxidation (FAO) pathway. While ω-oxidation has been shown to occur with monocarboxylic substrates as long as 16 carbons (i.e., palmitate), the preferred substrates are 10-12 carbons in length [21]. The metabolism of even-chain DCAs has been extensively investigated by our group and others [22–28]. In contrast, other than glutaric acid (hereafter, DC_5_), very little is known about odd-chain DCA metabolism. Their existence in the human body is evidenced by the detection of the seven (DC_7_, aka pimelic acid) and nine carbon (DC_9_, aka azelaic acid) species in urine and tissues. Accumulation of odd-chain DCAs up to 15 carbons in length are commonly observed in urine and blood from patients with inborn errors of metabolism, most notably peroxisomal disorders [29–31]. The origin of these longer odd-chain DCAs has remained enigmatic, as has their metabolic fates. Here, we hypothesized that the β-oxidation of longer odd-chain DCAs produces DC_5_, contributing to the substrate load on GCDH. This additional source of DC_5_, independent of lysine catabolism, could exacerbate symptoms in GA1, particularly during catabolic states in which the ω-oxidation pathway is strongly induced [21, 32–34].

## RESULTS

### Feeding DC_11_ biochemically phenocopies glutaric aciduria type 1 in mice

In our previous work, we fed mice DC_12_ and observed significant increases in the 6-carbon dicarboxylic intermediates adipic acid (DC_6_) and adipoylcarnitine (DC_6_-carnitine) [26, 35]. Here, we hypothesized that feeding the odd-chain dicarboxylic DC_11_ would produce the odd-chain e intermediates glutaric acid (DC_5_) and glutarylcarnitine (DC_5_-carnitine), which are major diagnostic biomarkers for GA1 in urine and serum, respectively (Fig. 1a). Indeed, after 10 days of feeding 10% w/w DC_11_ to male 129S1 mice, we observed a marked increase in urinary odd-chain DCAs including DC_11_, DC_9_, DC_7_ and DC_5_ (Fig. 1b). In addition, we observed significant excretion of unsaturated and 3-hydroxy odd-chain DCAs including 3-hydroxyundecanedioic (3OH-DC_11_), 3-hydroxyazelaic (3OH-DC_9_), 3-hydroxypimelic acid (3OH-DC_7_) and 2-enoyl-DC_7_ (Fig. S1). In general, the urinary concentration of these metabolites increased with decreasing chain length, with DC_5_ being the most abundant. Interestingly, in liver tissue harvested from these same animals the pattern was different. DC_5_ was highly abundant in control-fed liver and did not change with DC_11_ feeding (Fig. 1c). Rather, the most prominent change associated with DC_11_ diet was the presence of a large amount of DC_7_, which was undetectable in control-fed liver. The liver may be adapted to either catabolize or excrete glutaric acid to prevent its accumulation. In serum, DC_5_-carnitine was increased ∼10-fold (Fig. 1d).

**Figure 1.**
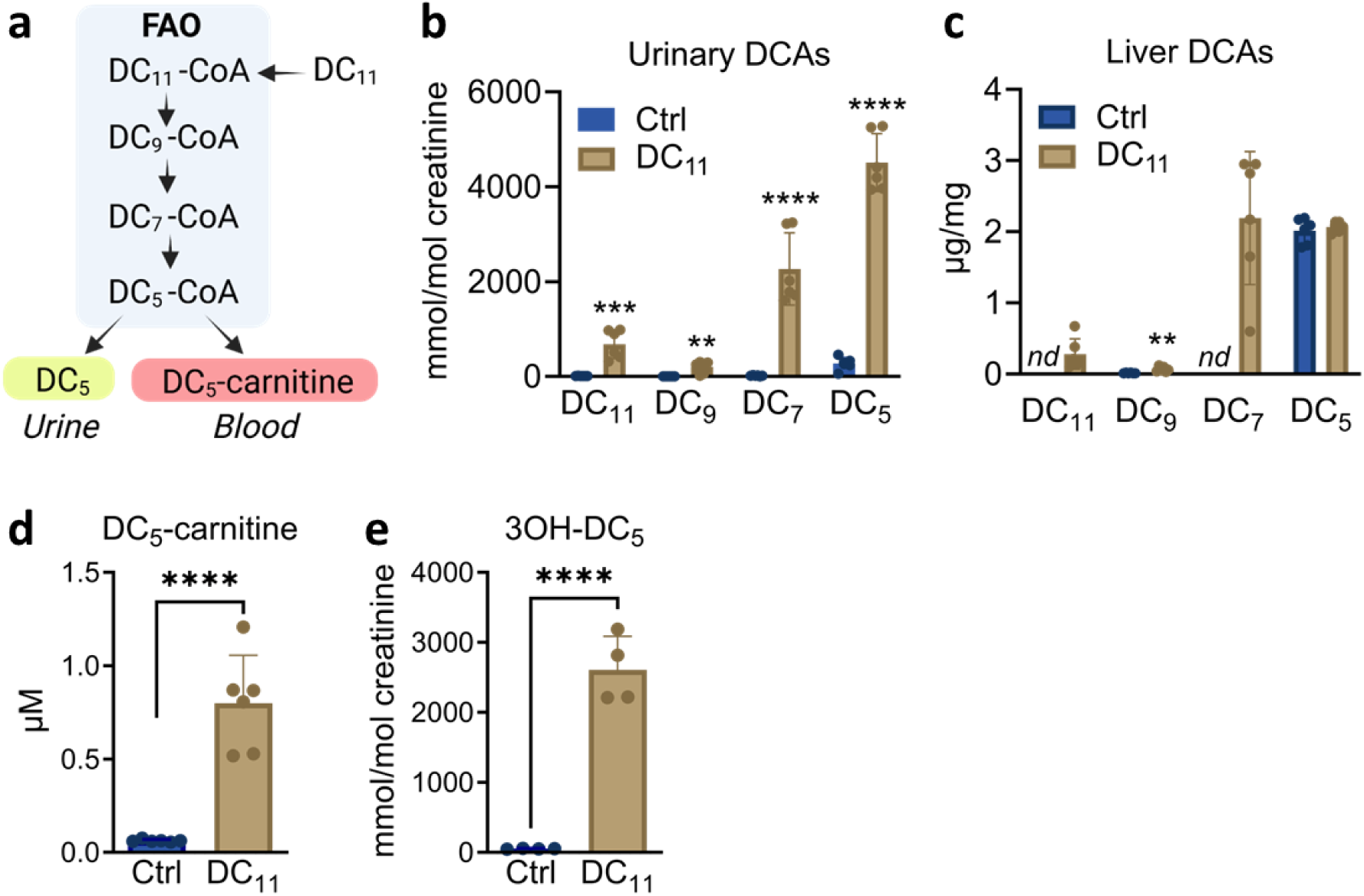
Feeding DC_11_ to wildtype mice recapitulates the biochemical phenotype of glutaric aciduria type 1 (GA1). **a)** It is hypothesized that chain-shortening DC_11_ produces DC_5_ and its acylcarnitine conjugate, DC_5_-carnitine, which both exit the cell. b) Male 129S1 mice were fed DC_11_ at 10% w/w for 10 days and urine used for mass spectrometry. The starting compound DC_11_ and sequentially chain-shortened intermediates were significantly increased in urine. c) Liver tissue from the same animals in (b) was also subjected to mass spectrometry. d) DC_5_-carnitine was measured in serum by mass spectrometry. e) Mass spectrometry detected the GA1 metabolite 3-hydroxyglutaric acid (3OH-DC^5^) in urine after 10 days on DC_11_ diet. All urinary analyses were normalized to creatinine and liver analyses to tissue weight; *nd* = not dectected. All graphs represent means and standard deviations. Student’s t-test was used to compare groups. **P<0.01, ***P<0.001, ***P<0.0001.

In addition to urinary DC_5_ and serum DC_5_-carnitine, most GA1 patients also present with significant urinary excretion of glutaconic acid (2-enoyl-DC_5_) and 3-hydroxyglutaric acid (3OH-DC_5_), a metabolite considered pathognomonic for GA1 [17, 36, 37]. Surprisingly, we observed that consumption of DC_11_ produced significant increases in urinary 3OH-DC_5_ (Fig 1e) as well as glutaconic acid. Thus, DC_11_ consumption in mice recapitulates the biochemical profile associated with GA1.

### DC_11_ induces protein glutarylation across multiple cellular compartments in the liver

GCDH is a mitochondrial enzyme, and the diagnostic metabolites associated with GA1 (DC_5_, 3OH-DC_5_, and DC_5_-carnitine) originate in mitochondria. In contrast, DCAs produced by omega oxidation in liver and kidney are thought to be degraded through peroxisomal FAO [21]. To better understand how DC_11_ consumption recapitulates the biochemical phenotype of GA1, we next sought to interrogate the compartmentalization of DC_11_ chain-shortening using protein glutarylation as a biomarker for glutaryl-CoA abundance. Among acyl-CoA metabolites, the dicarboxylic species DC_4_-CoA (succinyl-CoA) and DC_5_-CoA are known to exhibit high chemical reactivity toward the lysine residues of proteins [38, 39]. We previously used protein succinylation to follow the metabolism of DC_12_ to DC_4_-CoA [26, 35]. Here, we used the same strategy, measuring protein glutarylation as a surrogate for DC_5_-CoA. We began by immunoblotting mouse tissues with a pan anti-glutaryllysine antibody, using tissues harvested from mice fed DC_11_ for 10 days. Muscle, heart, and brain showed no change in protein glutarylation with DC_11_ feeding (data not shown), while liver and kidney both showed dramatic protein hyperglutarylation (Fig. 2a). Because liver and kidney are rich in peroxisomes, these immunoblots suggested that DC_11_ may be chain-shortened to DC_5_-CoA through peroxisomal β-oxidation rather than mitochondrial β-oxidation. In keeping with this, lysates prepared from purified hepatic peroxisomes exhibited a strong reactivity with anti-glutaryllysine antibody in DC_11_-fed mice (Fig. 2b). However, mass spectrometry profiling of lysine glutarylation in DC_11_-fed liver tissue revealed that DC_11_ induces protein glutarylation broadly across multiple compartments, including both peroxisomes and mitochondria. A total of 1,823 glutarylated peptides were quantified in the analysis, representing 643 proteins. DC_11_ feeding was associated with a statistically significant hyper-glutarylation of 83% of all glutarylated peptides (Fig. 2c, Supplemental Table 1). The median Log2 fold-change in lysine glutarylation across these peptides was 9.5 (724-fold). To visualize the subcellular distribution of these changes we utilized a method known as localization of organelle proteins by isotope tagging (LOPIT) [40]. Fig. 2d shows an overlay of the glutarylation data onto the LOPIT reference map created by the Lilley group ([41]; see Fig. S2), with red color indicating increased glutarylation and the size of the spots indicating the log2 fold-change. DC_11_ induced strong hyperglutarylation most prominently in mitochondria, peroxisomes, endoplasmic reticulum, and cytoplasm, indicating the presence of DC_5_-CoA in all these compartments.

**Figure 2.**
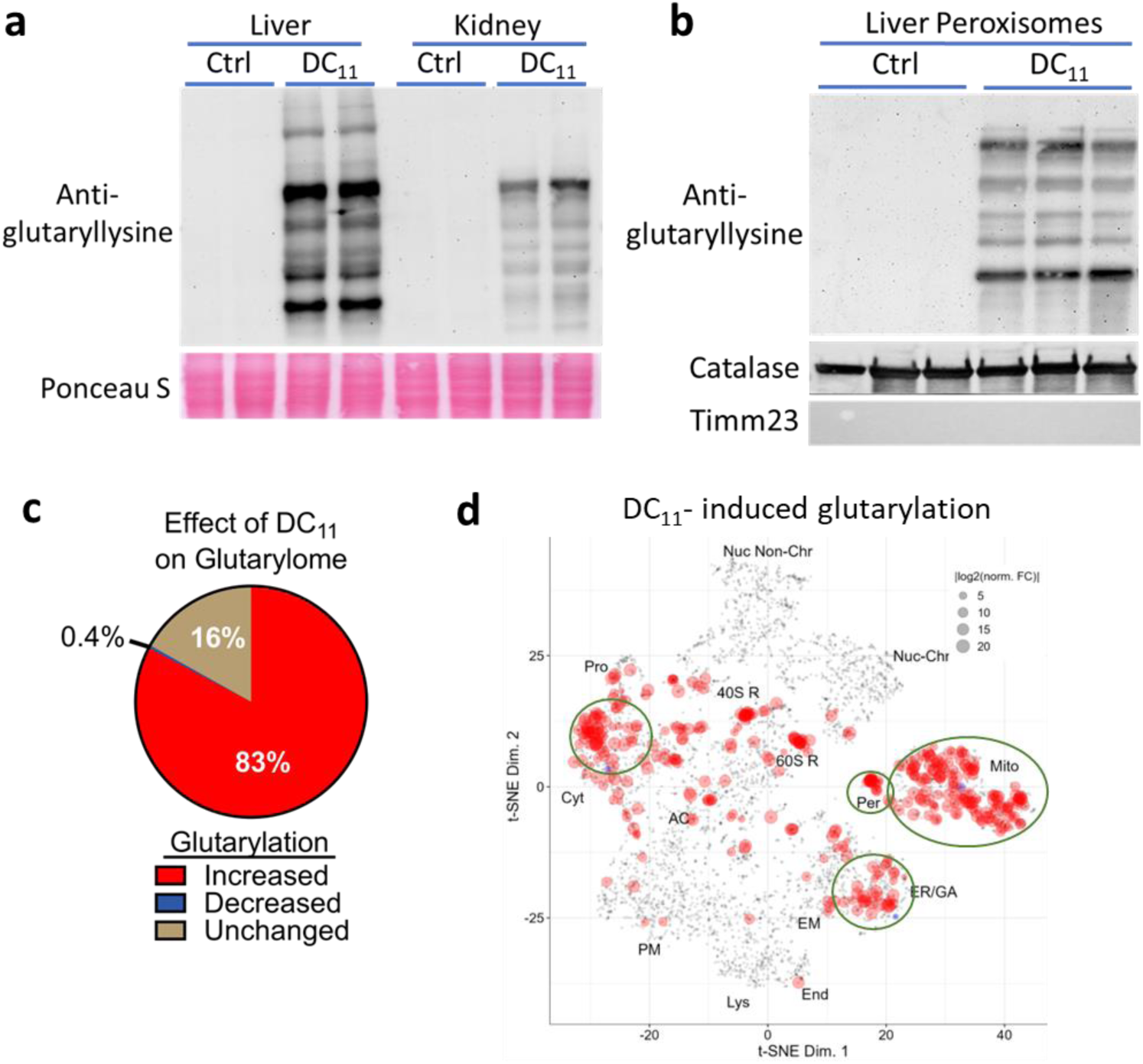
DC_11_ induces protein glutarylation across multiple cellular compartments in the liver. Protein glutarylation was followed as a proxy for glutaryl-CoA (DC_5_-CoA), which chemically reacts with lysine residues. a) Male mice were fed DC_11_ at 10% w/w for 10 days or standard diet (Ctrl). Liver and kidney lysates (25 µg) were immunoblotted with a pan anti-glutaryllysine antibody. Note that Ctrl tissues exhibit low signal that requires extended exposure to visualize. b) Liver peroxisomes were purified following Ctrl or DC_11_ diet and immunoblotted with anti-glutaryllysine antibody (10 µg per lane). Catalase was used as a peroxisomal marker and Timm23 a mitochondrial marker to demonstrate purity of the peroxisomal fraction. c,d) Mass spectrometry was used to profile the lysine glutarylome in N=5 male mice fed Ctrl or DC_11_ diets for 10 days. 1,823 glutarylated peptides were quantified, representing 643 unique proteins. Abundance of glutarylated peptides from a given protein was corrected for any inter-group differences in abundance of the whole protein. The majority (83%) of the 1,823 glutarylated peptides were significantly increased on DC_11_ diet (q<0.05, >1.5 fold-change). The average fold-change increase was 724-fold. Panel (d) shows a LOPIT plot distributing the glutarylation signal across intracellular compartments. The largest changes in glutarylation on DC_11_ diet occurred in four compartments circled in green on the plot: mitochondria, peroxisomes, endoplasmic reticulum/golgi apparatus, and cytoplasm. A reference map with full definitions of the compartments appears in Supplemental Figure 1.

### Peroxisomes are the primary catabolic site for DC_11_ in HEK293 cells

The hyper-glutarylation observed on hepatic mitochondrial proteins from DC_11_-fed mice indicates that DC_11_ feeding increases DC_5_-CoA levels inside mitochondria. One possible explanation would be direct β-oxidation of DC_11_ to DC_5_-CoA via the mitochondrial FAO pathway. To further investigate the compartmentalization of DC_11_ catabolism we used the HEK-293 cell line model, which we have previously demonstrated to metabolize DC_12_ and DC_16_ and produce chain shortened β-oxidation intermediates [24, 42]. To determine if mitochondria can contribute to DC_11_ metabolism directly, we eliminated peroxisomes from HEK-293 cells. PEX13 is a key peroxisomal biogenesis factor, and *PEX13* knockout (KO) HEK-293 cells have no detectable peroxisomes [42]. Wild-type parental HEK-293 cells and PEX13 knockout cells were incubated with DC_11_ and then analyzed for the presence of DC_11_ catabolic products. DC_11_ treatment of wild-type HEK-293 cells led to accumulation of DC_9_, DC_7_, and DC_5_ in the media and increased the abundance of their acylcarnitine conjugates in cells (Fig. 3). This was most pronounced for DC_9_ and DC_7_, which were undetectable in media of vehicle-treated wild-type HEK-293 cells (Fig. 3a,b). Interestingly, in the absence of peroxisomes, DC_11_ treatment was still able to increase the abundance of DC_5_, DC_5_-carnitine, DC_7_-carnitine, and DC_9_-carnitine by ∼2-fold (Fig. 3c-f). These data are consistent with peroxisomes being the primary site of DC_11_ catabolism and mitochondrial FAO playing a minor role in the absence of peroxisomes. Further, these data indicates that metabolites associated with GA1, normally thought to be strictly mitochondrial in origin, can come from the peroxisome when odd-chain DCAs are present.

**Figure 3.**
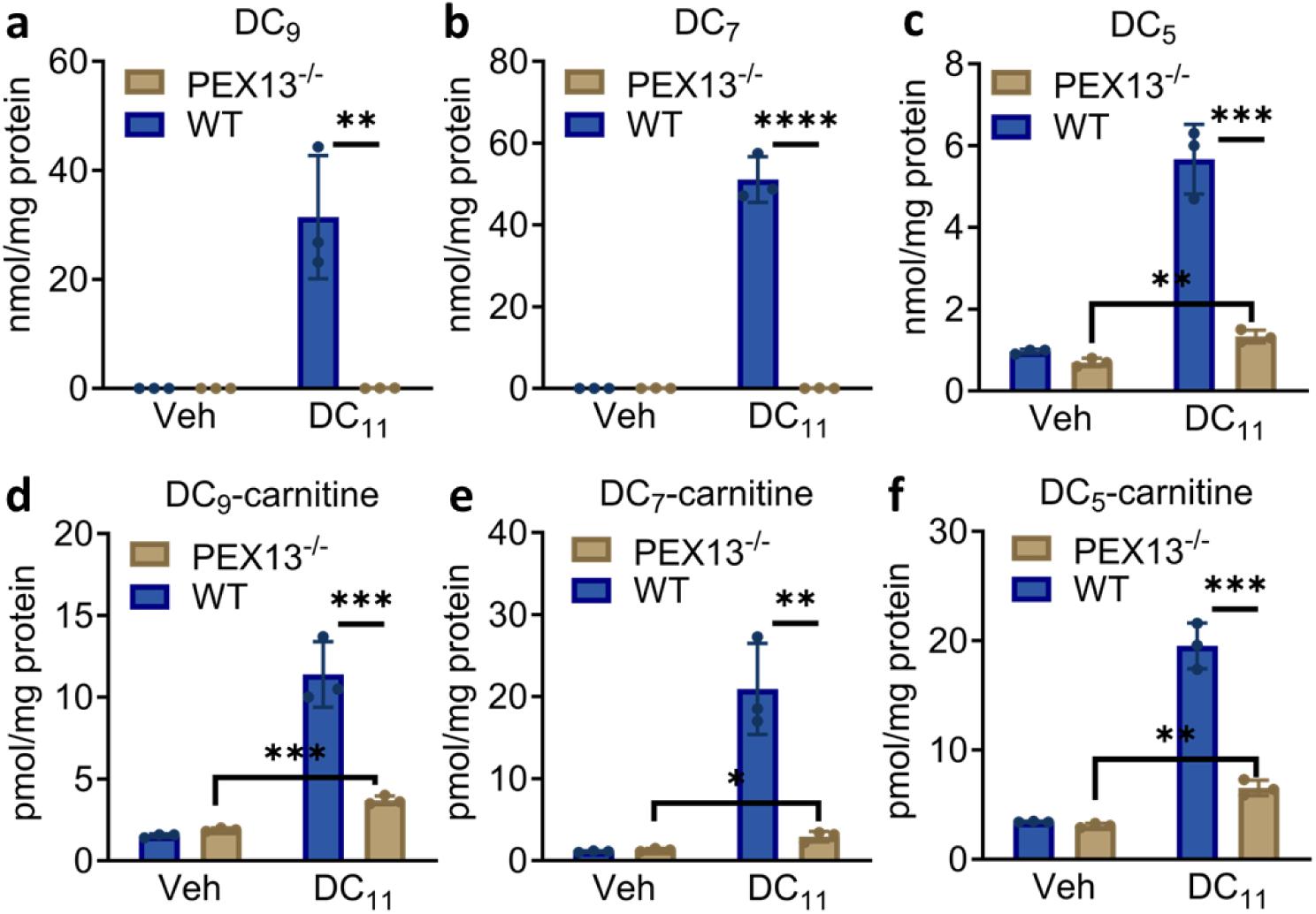
Peroxisomes are the primary catabolic site for DC_11_ in HEK-293 cells. Parental wild-type HEK-293 cells and HEK-293 cells modified by CRISPR/Cas9 to lack the peroxisomal biogenesis factor PEX13 were incubated with DC_11_ for 72 hours. Dicarboxylic acids were quantified in the media (a-c) and odd-chain dicarboxylic acylcarnitines in the cell pellet (d-f). All concentrations were normalized to cellular protein. All graphs represent means and standard deviations. Student’s t-test was used for pairwise comparisons of 1) PEX13 KO vehicle-treated *vs* PEX13 KO DC_11_-treated cells; and 2) wild-type DC_11_-treated *vs* PEX13 KO DC_11_-treated cells. *P<0.05, **P<0.01, ***P<0.001, ***P<0.0001.

### Succinyl-CoA:glutarate-CoA transferase is required for mitochondrial oxidation of externally generated DC_5_

It is widely held that acyl-CoAs are highly compartmentalized and cannot cross organelle membranes intact. Acyl-CoAs are either hydrolyzed by thioesterases to their free acids or esterified to carnitine by acyltransferases for transport across membranes [43]. Peroxisomal DC_5_ transferring to mitochondria must be re-esterified to CoA to be further metabolized by GCDH. We hypothesized that the poorly understood enzyme succinyl-CoA:glutarate-CoA transferase (SUGCT) may facilitate the mitochondrial metabolism of DC_5_ released by peroxisomes. SUGCT uses succinyl-CoA as a donor to generate DC_5_-CoA from free DC_5_ [44]. Interestingly, we previously demonstrated that 129S2/SvPasCrl are naturally deficient in the enzyme SUGCT, and further data suggested that this a biochemical trait common to other 129 substrains as well [45]. Coincidentally, the mice utilized here for DC_11_ feeding were 129S1/SvImJ. We then established that the 129S1 substrain is indeed deficient in SUGCT (Fig. 4a,b). Our observation that feeding DC_11_ could increase mitochondrial protein glutarylation in SUGCT-deficient mice suggests that SUGCT is not required for the intramitochondrial enrichment in DC_5_-CoA. To test this, we fed DC_11_ to a cohort of 129S1 mice and a cohort of C57BL/6 mice, which abundantly express SUGCT, and used anti-glutaryllysine immunoblotting of liver lysates to compare the profiles of global protein hyperglutarylation. The overall profiles were strikingly similar (Fig. 4c). To determine if other unidentified protein(s) could activate DC_5_ to DC_5_-CoA, we used ^14^C-DC_5_ and followed its metabolism to ^14^C-CO_2_. To yield ^14^C-labeled CO_2_, the ^14^C-DC_5_ would need to enter mitochondria, become activated to CoA, and then be decarboxylated by GCDH. Kidney and liver lysates from SUGCT-deficient (129S1) and SUGCT-expressing (C57BL/6) were assayed for their ability to generate ^14^C-CO_2_ from ^14^C-DC_5_. The loss of SUGCT reduced DC_5_ oxidation rates by 90% in liver homogenates and 95% in kidney homogenates, confirming that the primary route for catabolizing free DC_5_ in mitochondria is through SUGCT (Fig 4d). This experiment suggests that 129S1 mice retain a small amount of residual SUGCT activity (5-10%) that is sufficient to mediate the protein glutarylation phenotype. A role for SUGCT in DC_11_ catabolism was further demonstrated by profiling urinary odd-chain DCAs in C57BL/6 and 129S1 mice following DC_11_ feeding. The amount of excreted DC_9_ and DC_7_ was equal between strains but the amount of excreted DC_5_ was 60% lower in C57BL/6 mice, consistent with processing of DC_5_ through SUGCT (Fig. 4e).

**Figure 4.**
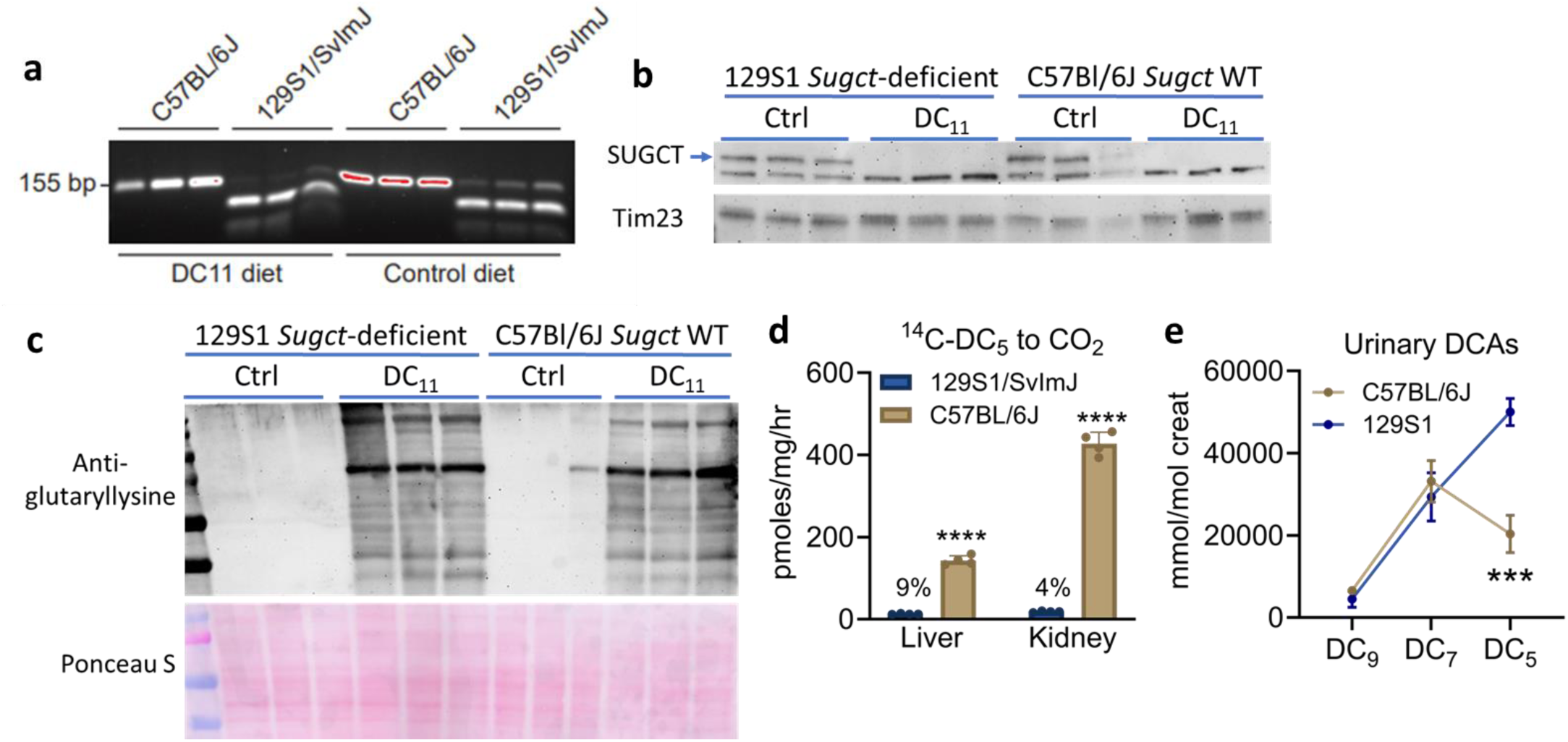
Succinyl-CoA:glutarate-CoA transferase is required for mitochondrial oxidation of externally generated DC_5_. **a,b)** 129S1/SvImJ (129S1) mice carry a naturally-occurring mutation in the *Sugct* gene as evidenced by PCR genotyping (a) and immunoblotting for SUGCT protein (b). c) Male 129S1 mice lacking SUGCT and C57BL/6J mice wildtype for SUGCT were fed standard diet (Ctrl) or 10/% w/w DC_11_ diet for 10 days. Liver lysates (25 µg) were immunoblotted with a pan anti-glutaryllysine antibody. DC_11_ has approximately equal effects on glutarylation across mouse strains. d) Liver and kidney homogenates from 129S1 mice lacking SUGCT and C57BL/6J mice wild-type for SUGCT were used to measure mitochondrial oxidation of ^14^C-DC_5_ to ^14^C-CO_2_. Loss of SUGCT compromises the ability to activate DC_5_ to CoA for oxidation by GCDH. e) 129S1 mice lacking SUGCT and C57BL/6J mice wild-type for SUGCT were fed 10/% w/w DC_11_ for 10 days and urine was subjected to mass spectrometry for odd-chain dicarboxylic acids. Graphs represent means and standard deviations. Student’s t-test was used for pairwise comparisons of 129S1 to C57BL/6 mice. ***P<0.001, ***P<0.0001.

### Peroxisomes may generate the GA1 biomarker 3OH-DC_5_ during β-oxidation of DC_5_ to DC_3_

The experiments in Figure 3 demonstrate that DC_11_ catabolism can produce the GA1 metabolites DC_5_ and DC_5_-carnitine inside peroxisomes. This raises the question of whether peroxisomes can also produce 3OH-DC_5_, the pathognomonic marker of GCDH deficiency, which we showed is also elevated in urine of mice fed DC_11_, despite being wild-type for GCDH (Fig. 1e). While SUGCT deficiency reduced the amount of urinary DC_5_ (Fig. 4e), it did not alter excretion of 3OH-DC_5_ (Fig. 5a). This suggests that the 3OH-DC_5_ that accumulates with DC_11_ consumption is uncoupled from mitochondrial metabolism of DC_5_. In mitochondria, when GCDH is absent the FAO machinery attempts to chain-shorten DC_5_-CoA to DC_3_-CoA (malonyl-CoA). The DC_5_-CoA is dehydrogenated by MCAD and then hydrated by 3MGH to produce 3OH-DC_5_, but there are no enzymes capable of catalyzing the final two steps to convert 3OH-DC_5_-CoA to DC_3_-CoA and therefore the 3OH-DC_5_-CoA accumulates. After hydrolysis it is released as the free 3OH-DC_5_ acid. The peroxisomal FAO machinery accommodates a very broad range of substrates [46, 47]. Historically, peroxisomal FAO was thought to shorten fatty acids to eight carbons and then release the medium-chain products to the mitochondria for completion. However, in recent years it has been shown that peroxisomal FAO can chain-shorten even-chain substrates two more rounds, albeit inefficiently, to yield four-carbon products [26, 48, 49]. Odd-chain substrates have not been explored. Here, we show that peroxisomal FAO can shorten DC_5_-CoA to DC_3_-CoA, and therefore, can produce the GA1 biomarker 3OH-DC_5_ as an FAO intermediate. First, we observed a significant increase in DC_3_ (malonic acid) in the urine of wild-type mice fed DC_11_ (Fig. 5b). To put this into context with upstream metabolites, we compared the urinary concentration of DC_3_ to that of DC_5_ and the intermediate 3OH-DC_5_ (Fig. 5c). The amount of 3OH-DC_5_ and DC_3_ were about 12% and 4% that of DC_5_, respectively, consistent with a slow, inefficient oxidation of DC_5_-CoA to DC_3_-CoA. We next assayed the activity of recombinant acyl-CoA oxidase-1a (ACOX1a), the rate-limiting enzyme for DCA oxidation in peroxisomes, with DC_5_-CoA as substrate. ACOX1a exhibited activity with DC_5_-CoA with an approximate *K*m of 42 µM, which about 2X higher than the *K*m we previously reported for ACOX1a with DC_6_-CoA and nearly 10X higher than the *K*m with DC_12_-CoA [26]. Therefore, analogous to the known production of DC_4_-CoA (succinyl-CoA) from even-chain DCAs, peroxisomes may produce DC_3_-CoA (malonyl-CoA) from odd-chain DCAs.

**Figure 5.**
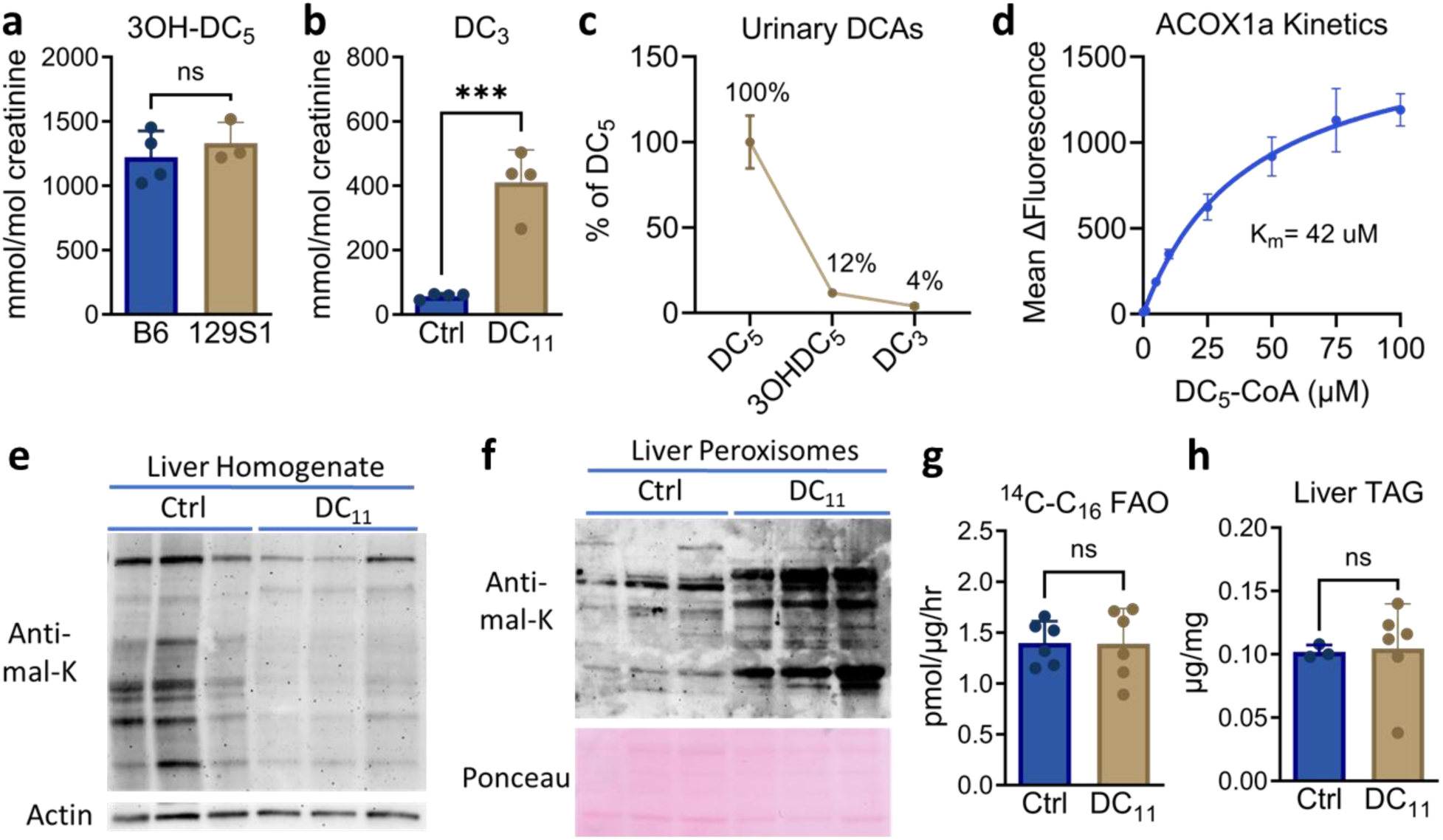
Peroxisomes may generate the GA1 biomarker 3OH-DC_5_ during β-oxidation of DC_5_ to DC_3_. **a)** 129S1 mice lacking SUGCT and C57BL/6J mice wildtype for SUGCT were fed 10/% w/w DC_11_ for 10 days and mass spectrometry used to measure 3OH-DC_5_ in urine. a) Urine from C57BL/6J mice fed DC_11_ was assayed for malonic acid (DC_3_) by mass spectrometry. c) The urinary concentrations of 3OH-DC_5_ and DC_3_ were normalized to that of DC_5_ in male C57BL/6J mice fed DC_11_. d) Purified recombinant ACOX1a demonstrates activity with DC_5_-CoA as substrate with an estimated *K_m_* of 42 µM. e) Male 129S1 mice were fed standard diet (Ctrl) or 10/% w/w DC_11_ diet for 10 days. Liver lysates (25 µg) were immunoblotted with a pan anti-malonyllysine (mal-K) antibody. f) Liver peroxisomes were purified following Ctrl or DC_11_ diet and immunoblotted with anti-malonyllysine antibody (10 µg per lane). g,h) Liver from male 129S1 mice fed standard diet (Ctrl) or 10/% w/w DC_11_ diet for 10 days was used to measure mitochondrial fatty acid oxidation (FAO) with ^14^C-labeled palmitate (C_16_) and triglyceride content. Graphs represent means and standard deviations. Student’s t-test was used for pairwise comparisons. ***P<0.001.

Although weaker than DC_5_-CoA or DC_4_-CoA, DC_3_-CoA is also reactive with lysine residues on the surface of proteins [39]. We used anti-malonyllysine immunoblotting to probe the global levels of protein malonylation in liver homogenates from mice fed control diet or DC_11_. Surprisingly, the global malonylation levels were lower in DC_11_-fed livers (Fig. 5d), which may be related to the extensive glutarylation (Fig. 2). However, when isolated peroxisomal protein was analyzed, there was a clear increase in malonylation (Fig. 5e). The discrepancy between the reduced global protein malonylation and the higher peroxisomal protein malonylation suggests that, unlike what we observed for DC_5_-CoA, the DC_3_-CoA formed inside peroxisomes does not give rise to DC_3_-CoA in other compartments. In the cytoplasm, DC_3_-CoA promotes fatty acid synthesis while inhibiting mitochondrial FAO via inhibition of carnitine palmitoyltransferase-1. We measured the rate of mitochondrial FAO in DC_11_-fed liver homogenates with the CPT1 substrate ^14^C-palmitate and observed no effect of DC_11_ diet on FAO (Fig. 5g). Similarly, the amount of liver triglyceride was unchanged after 10 days on DC_11_ diet (Fig. 5h). These data suggest that DC_3_-CoA produced in the peroxisome is not altering fatty acid homeostasis in other compartments.

## DISCUSSION

In these studies, we have shown that β-oxidation of odd-chain DCAs can reproduce the biochemical phenotype of GA1, an inborn error of lysine metabolism. Our data is consistent with a model in which DC_11_ is primarily oxidized by peroxisomes in the liver, and likely to a lesser extent in kidney (Fig. 6). Three rounds of chain-shortening through β-oxidation produces DC_5_-CoA, which then has three fates: 1) thioesterase cleavage to free DC_5_, likely by the enzyme acyl-CoA thioesterase-8 (ACOT8) [48, 50]; 2) esterification to carnitine, likely by the enzyme carnitine octanoyltransferase (CROT) [51]; or 3) conversion to DC_3_-CoA by one more round of β-oxidation, producing the GA1 diagnostic hallmark metabolite 3OH-DC_5_ in the process. Peroxisomes are known to harbor malonyl-CoA decarboxylase [52], which converts DC_3_-CoA into acetyl-CoA. All these products can leave the peroxisome for further cellular metabolism, for example in the mitochondria. They can also end up in circulation and ultimately in the urine.

**Figure 6.**
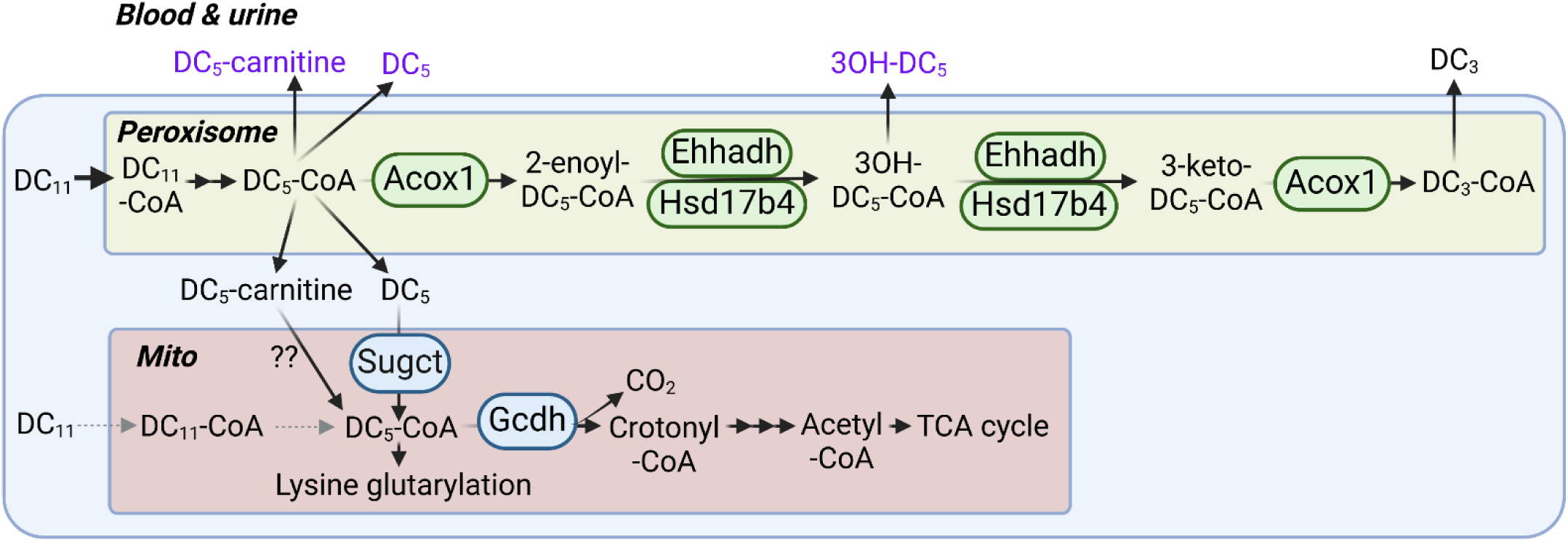
Proposed model for generation of a GA1-like biochemical signature during odd-chain dicarboxylic acid metabolism. DC11 is primarily oxidized by peroxisomal FAO in liver. Peroxisomes chain-shorten DC11-CoA to DC5-CoA with three rounds of β-oxidation. DC5-CoA has three fates: 1) release from peroxisomes as free DC5 after thioesterase cleavage; 2) release from peroxisomes as DC5-carnitine after CoA/carnitine exchange by a carnitine acyltransferase enzyme; or 3) further, slow oxidation to DC3-CoA. The GA1 signature metabolite 3OH-DC5 is formed as an intermediate in the conversion of DC5-CoA to DC3-CoA. DC5 released by peroxisomes can be oxidized by mitochondria, thereby increasing the substrate load on the enzymes SUGCT and GCDH, which are mutated in glutaric aciduria type 3 and glutaric aciduria type 1, respectively.

DC_5_ can be re-activated to CoA inside the mitochondria by SUGCT, producing DC_5_-CoA which is then metabolized by GCDH and downstream enzymes. The importance of SUGCT for eliminating DC_5_ produced in peroxisomes is highlighted by the 60% higher urinary excretion of DC_5_ in 129 mice, which have a naturally occurring deficiency of SUGCT. SUGCT is the enzyme deficient in GA3. GA3 is characterized by glutaric aciduria in the absence of increased 3-OH-DC_5_ and DC_5_-carnitine [53]. Based on our results it is possible that the DC_5_ accumulating in GA3 is at least partly of peroxisomal origin. We did not directly investigate the fate of peroxisomal DC_5_-carnitine in these studies. However, mitochondria are well known to interchange DC_5_-CoA and DC_5_-carnitine, and we postulate that DC_5_-carnitine released from peroxisomes could also contribute to the intramitochondrial pool of DC_5_-CoA, possibly by the action of carnitine palmitoyltransferase-2 (CPT2).

Another remarkable finding was the identification of 3OH-DC_5_, considered pathognomonic for GA1, in urine of DC_11_ fed mice. Prior work has revealed that even-chain dicarboxylic aciduria during fasting or in patients with mitochondrial FAO defects is accompanied by excretion 3-hydroxy even-chain intermediates including 3-OH-DC_6_ and its cyclic lactone form [54, 55]. These metabolites likely represent the metabolic intermediates in the peroxisomal β-oxidation of DC_6_ to DC_4_. Upon feeding of DC_11_, we similarly observed excretion of 3-hydroxy and unsaturated odd-chain intermediates, again likely reflecting peroxisomal β-oxidation of DC_11_. Based on our data, we propose that the 3OH-DC_5_ observed in the urine of DC_11_ fed mice has a peroxisomal origin, which differs from the mitochondrial origin in GA1. Importantly, it is generally accepted that the acute encephalopathic crises in GA1 are caused by local cerebral generation and entrapment of toxic metabolites resulting from lysine degradation, such as DC_5_ and 3OH-DC_5_ [19, 20]. One recent study has challenged this notion through demonstrating that the toxic metabolites are produced by the liver and thus are able to cross the blood-brain barrier to exert their detrimental effect [56]. The latter mechanism would implicate odd-chain DCA metabolism in liver and kidney peroxisomes as an unrecognized source of toxic metabolites in GA1. Although lysine degradation is the major source of DC_5_ in GA1, the relative contribution of all metabolites contributing to the pool of accumulating of GCDH substrate, including hydroxylysine, tryptophan and odd-chain DCAs, will need to be determined in future studies.

Our studies have provided an initial exploration of odd-chain DCA metabolism and established that peroxisomal β-oxidation is a likely source of endogenous DC_9_, DC_7_ and DC_5_. Dietary sources of DCAs, however, are limited and therefore our mouse model of DC_11_ feeding provided supraphysiological amounts of odd-chain DCAs. The only known food containing DCAs is bran, which contains trace amounts (4.3 µmoles/g) of DC_9_ [57]. DC_9_ is also used as a topical treatment for mild to moderate acne, but the absorbed dose is low. But there are several potential endogenous sources of odd-chain DCAs including lipid peroxidation. Lipid peroxidation of 18-carbon fatty acids produces DC_9_ [58, 59]. In keeping with this, we have observed DC_9_ in liver, kidney, brain, heart, and muscle from wild-type mice maintained on a normal diet (data not shown). Another endogenous source could be ω-oxidation of monocarboxylic odd-chain fatty acids. Odd-chain fatty acids are readily detected in serum and tissue of rodents and humans and have both dietary and endogenous sources. There is considerable evidence that α-oxidation of even-chain fatty acids, such as the 2OH-fatty acids derived from the degradation of sphingolipids, are a significant source of odd-chain fatty acids [60]. How all these pathways ultimately contribute to the production of odd-chain DCAs remains to be addressed in future studies.

## METHODS

### Animals, DC_11_ administration, and Sugct genotyping

All animal protocols were approved by the University of Pittsburgh Institutional Animal Care and Use Committee (IACUC), and all experiments were conducted in accordance with the guidelines and regulations set forth in the Animal Welfare Act (AWA) and PHS Policy on Humane Care and Use of Laboratory Animals. 129S1 and C57BL/6J male mice aged 8 weeks were purchased from Jackson Laboratories (Bar Harbor, ME). The mice were maintained on a 12 hr light/dark cycle in a pathogen-free barrier facility. The DC_11_ diet was made by mixing DC_11_ free acid (Toronto Research Chemicals) by weight (10% w/w) into powdered standard rodent diet. The diet was provided in glass feeding jars for 10 days. Control animals received powdered diet without DC_11_. The *Sugct* locus was genotyped as described [45].

### HEK-293 cell line models

For the measurement of DCA metabolism, HEK-293 cells were seeded in 6-well plates (0.85 × 106 cells per well) and grown for 48 hours at 37°C in a humidified CO_2_ incubator (5% CO2, 95% air). Then the media were removed and 2 mL of fresh media were added to each well consisting of DMEM with 10% FBS, antibiotics and 400 µM L-carnitine further supplemented with 500 µM DC_11_ (added from a 50 mM stock in 100% ethanol) or vehicle. After incubation for 72h, the media were collected for organic acid analysis. The intracellular metabolites were immediately extracted by adding 1 mL of ice-cold extraction solvent (5 methanol:3 acetonitrile:2 water) directly to the plate with cells. Plates were immediately placed on dry ice and incubating overnight at −80 °C to aid protein precipitation. The following day, the cells were scraped, and the extracts were centrifuged at 20,000 × g for 20 min at 4°C. Supernatants (800 µL) were used for acylcarnitine analysis. One separate well was used to lyse cells in RIPA buffer for measurement of protein content using the bicinchoninic acid (BCA) assay with human serum albumin as calibrator.

### Metabolite analyses

Metabolite analyses shown in Figure 1b,c,d were performed by the University of Pittsburgh Health Sciences Mass Spectrometry Core. The remaining analyses were performed by the Mount Sinai Biochemical Genetic Testing Lab. Liver organic acids were normalized to tissue weight, while all urine data were normalized to creatinine. HEK-293 cell analyses were normalized to cellular protein. Urine was collected in metabolic caging over an 8-hr period during dark cycle (1:00 a.m. to 9:00 a.m.). Liver and serum were collected early in the light cycle. All organic acids were quantified using standard curves.

### Peroxisome purification

Peroxisomes were separated from freshly isolated liver tissue by density gradient centrifugation exactly as described previously [35, 61]. Briefly, ∼0.5 grams of minced tissue was homogenized with 10 slow up and down strokes in 5 mL of ice-cold Extraction Buffer (5 mM MOPS, pH 7.65, with 250 mM sucrose, 1 mM EDTA, 0.5% ethanol and 0.2 mM PMSF). The homogenate was cleared by sequential centrifugation: 1) 1000 × g for 10 minutes; 2) 2000 × g for 10 minutes; and 3) 25,000 × g for 20 minutes. The pellet was resuspended in Extraction Buffer and diluted with OptiPrep Density Gradient Medium (60% iodixanol, Sigma, D1556) and OptiPrep Dilution Buffer (5 mM MOPS, pH 8.0, with 1 mM EDTA and 0.1% ethanol) to make a diluted sample containing 22.5% OptiPrep. Samples were layered in ultracentrifuge tubes and centrifuged for 3 hours at 50,000 × g. The loose pellet at the bottom of the tube was taken as the pure peroxisome fraction. We have previously validated this method using immunoblotting for organelle markers [35, 61].

### Immunoblotting

Tissue lysates were electrophoresed on Criterion SDS polyacrylamide gels (Bio-Rad, Hercules, CA) and transferred to nitrocellulose membranes. Antibodies used were: anti-glutaryllysine and anti-malonyllysine (PTM Biolabs); anti-SUGCT (Proteintech); anti-actin (Sigma); anti-catalase (Sigma #C0979); and anti-mitochondrial import inner membrane translocase subunit TIM23 (BD Biosciences #611222). After incubation with HRP-conjugated secondary antibodies, blots were visualized with chemiluminescence (Bio-Rad Clarity family ECL reagents).

### Analysis of site-level protein glutarylation by mass spectrometry

Pieces of frozen liver (∼100 mg) were homogenized, alkylated, trypsinized, clarified, and desalted exactly as described [26]. 100 μg of each peptide elution was set aside for analysis of protein-level changes, and the remaining material used for enrichment of glutarylated peptides with the PTMScan glutaryl-Lysine Motif Kit (Cell Signaling Technologies). LC-MS/MS analyses were performed on a Dionex UltiMate 3000 system online coupled to an Orbitrap Eclipse Tribrid mass spectrometer (Thermo Fisher Scientific, San Jose, CA). All samples were acquired in data-independent acquisition (DIA) mode. DIA data was processed in Spectronaut (version 14.10.201222.47784) using directDIA for both the protein level as well as PTM enriched samples. All PTM results were corrected for any changes in protein expression detected by the protein-level analysis. The PTM data were then analyzed using a paired t-test and p-values were corrected for multiple testing. Significance was established as q-value < 0.05 and absolute Log2(fold-change) > 0.58. The dataset was then subjected to LOPIT analysis to visualize the subcellular distribution of glutarylated peptides. LOPIT was conducted as previously described [62]. Briefly, the hyperLOPIT2015 pluripotent stem cell dataset was downloaded in Rstudio using the pRolocData R package (Bioconductor) and plotted with the pRoloc R package. The t-SNE machine learning algorithm was used to reduce the number of dimensions in the LOPIT proteomic data to a 2D map where proteins cluster by similarity from multiple experimental factors. All LOPIT protein assignments were used as originally determined and without additional refinement.

### ^14^C substrate flux assays

1-^14^C-labeled palmitic acid (C_16_) was sourced from PerkinElmer (Waltham, MA) and ^14^C-DC_5_ from American Radiolabeled Chemicals (St. Louis, MO). Just prior to assay, the substrates were dried to completion under nitrogen. The C16 was solubilized in 10 mg/mL α-cyclodextrin via incubation at 37°C for 30 min while DC_5_ was solubilized in reaction buffer. The final substrate concentrations were 50 µM for C_16_ and 200 µM for DC_5_, containing 0.5 µCi per mL of ^14^C tracer. The reactions consisted of ∼100 µg of liver or kidney lysate in a volume of 200 µL containing reaction buffer (100 mM sucrose, 10 mM Tris-HCl, 5 mM KH_2_PO_4_, 0.2 mM EDTA, 80 mM KCl, 1 mM MgCl_2_, 0.1 mM malate, 0.05 mM coenzyme A, 2 mM ATP, and 1 mM DTT). The ^14^C-DC_5_ reactions also contained 1 mM citrate and 1 mM methylene blue. After 2 hr incubation at 37°C, the reactions were stopped by adding perchloric acid to 0.5 M final concentration. For ^14^C-DC_5_ reactions ^14^C-CO_2_ was captured on papers wetted with 2M KOH inserted into the tube caps. The ^14^C-C_16_ reactions were frozen overnight, centrifuged to remove precipitated material, and then the water-soluble ^14^C-labeled FAO products were isolated by methanol-chloroform extraction. All results were quantified on a scintillation counter and normalized to protein concentration.

### ACOX1 expression, purification, and activity assay

ACOX1a expression and purification have been described previously [26]. The human ACOX1a coding sequence with a 6xHis tag coding sequence at 5’-end was transformed into C43(DE3) competent cells and cultured at 37°C in LB medium. Expression was induced with 0.5 mM IPTG at 23°C for 3 hours. Cell pellets were lysed on ice by sonication, cleared by ultracentrifugation, and the recombinant protein was purified on a HisTrap HP column (Cytiva Sweden AB) using an AKTA Pure system. For activity, a constant amount (1 µg) of ACOX1 protein was assayed with a concentration range of DC_5-_ CoA by following H_2_O_2_ production with Amplex Red as described [25, 26].

