## Supplemental figures for "Odd-chain dicarboxylic acid feeding recapitulates the biochemical phenotype of glutaric aciduria type 1 in mice"

### SUPPLEMENTAL DATA

**Fig S1.** Urinary organic acid profile of a C57BL/6J mouse on $DC_{11}$ diet.

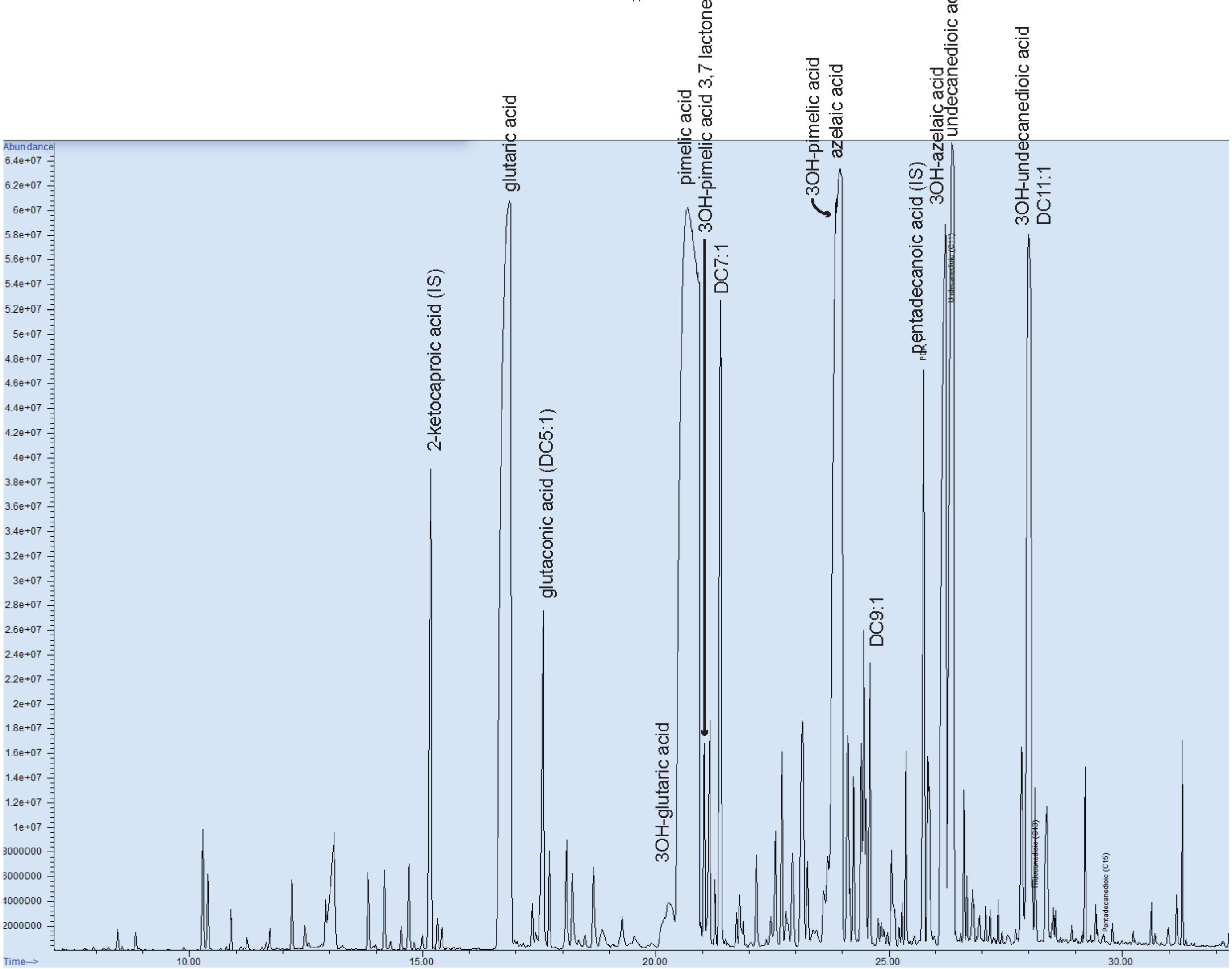


**Figure S1. Urinary organic acid profile of a C57BL/6J mouse on $DC_{11}$ diet.** The main peaks related to $DC_{11}$ metabolism are indicated as well as the internal standards (IS). $DC_{11}$, $DC_9$, $DC_7$ and $DC_5$ were quantified based on a dose response curve using authentic standards and a fixed amount of the pentadecanoic acid IS after dilution of urines to avoid peak saturation (20x). The 3OH odd-chain intermediates were identified using a m/z 233.1 fragment common for all 3-hydroxy carboxylic acids in combination with the specific $[M - 15]^+$ ion formed by loss of a methyl group bonded to the silicon atom (3OH $DC_{11}$ m/z 433.3, 3OH $DC_9$ m/z 405.2 and 3OH $DC_7$ m/z 377.2). The unsaturated dicarboxylic acids were identified based on the $[M - 15]^+$ ion ($DC_{11}$ m/z 343.2, $DC_9$ m/z 315.2, $DC_7$ m/z 287.2 and $DC_5$ m/z 259.1). A putative peak for 3OH $DC_7$-3,7 lactone was identified based on a previously reported mass spectrum of 3OH $DC_6$ 3,6-lactone ($[M - 15]^+$ = 215.1, the possible loss of mass fragments 28 (CO) from the molecular ion to form m/z 202.1 and the loss of mass fragment 44 ($CO_2$) from m/z 215.1 to yield m/z 171.1).

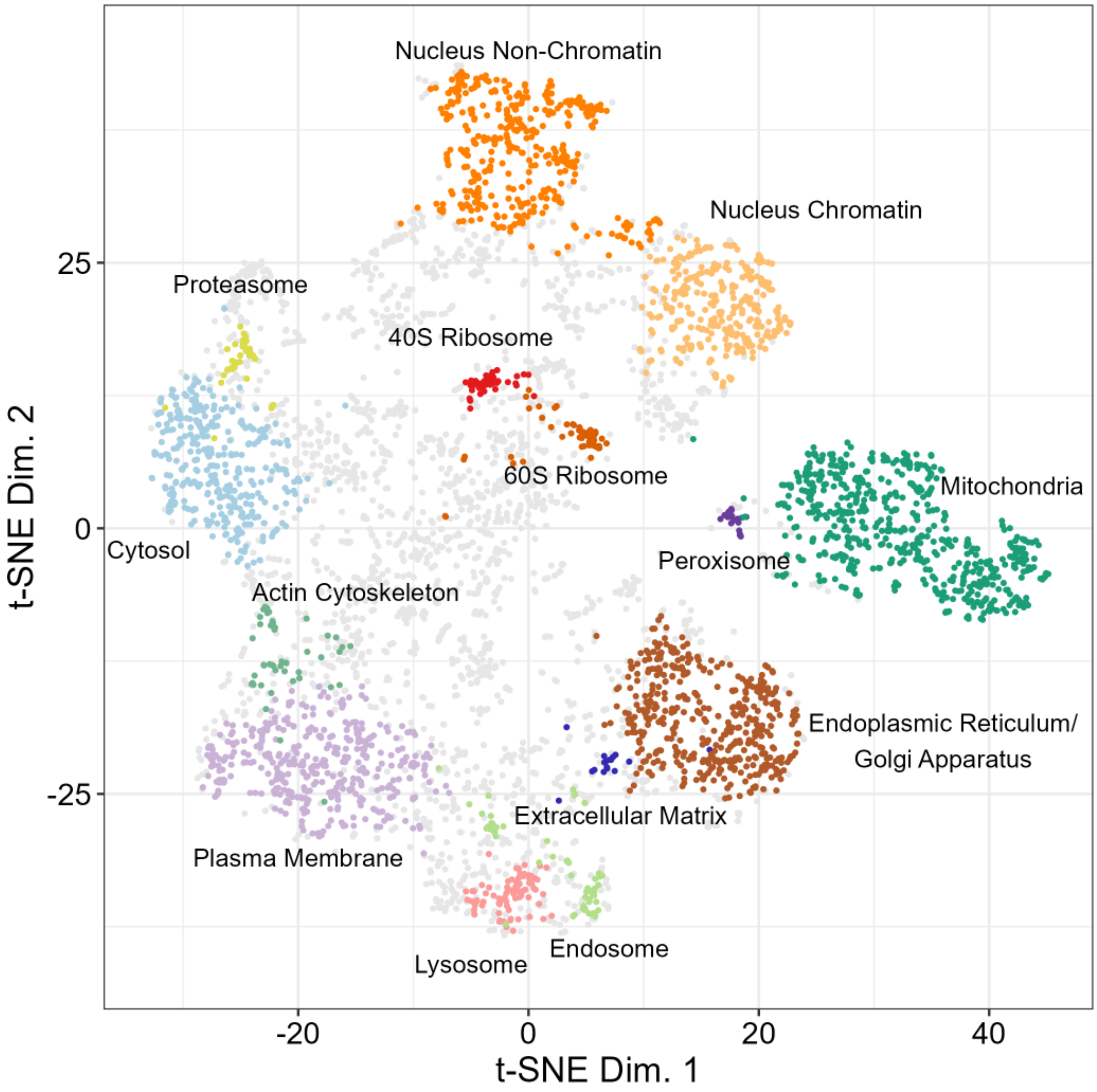


**Fig S2. Reference LOPIT map.** Distribution of 5,032 LOPIT-localized proteins from mouse pluripotent stem cells from Christoforou *et al.* This map was used to create the map of lysine glutarylation in Figure 2.
